# Self-supervised deep learning uncovers the semantic landscape of drug-induced latent mitochondrial phenotypes

**DOI:** 10.1101/2023.09.13.557636

**Authors:** Parth Natekar, Zichen Wang, Mehul Arora, Hiroyuki Hakozaki, Johannes Schöneberg

**Author notes:** Address correspondence to: Johannes Schöneberg, Ph.D., Departments of Pharmacology and Chemistry and Biochemistry University of California, San Diego, Center for Neural Circuits and Behavior, Room 106B, La Jolla, CA 92093.

## Abstract

Imaging-based high-content screening aims to identify substances that modulate cellular phenotypes. Traditional approaches screen compounds for their ability to shift disease phenotypes toward healthy phenotypes, but these end point-based screens lack an atlas-like mapping between phenotype and cell state that covers the full spectrum of possible phenotypic responses. In this study, we present MitoSpace: a novel mitochondrial phenotypic atlas that leverages self-supervised deep learning to create a semantically meaningful latent space from images without relying on any data labels for training. Our approach employs a dataset of ∼100,000 microscopy images of Cal27 and HeLa cells treated with 25 drugs affecting mitochondria, but can be generalized to any cell type, cell organelle, or drug library with no changes to the methodology. We demonstrate how MitoSpace enhances our understanding of the range of mitochondrial phenotypes induced by pharmacological interventions. We find that i) self-supervised learning can automatically uncover the semantic landscape of drug induced latent mitochondrial phenotypes and can map individual cells to the correct functional area of the drug they are treated with, ii) the traditional classification of mitochondrial morphology along a fragmented to fused axis is more complex than previously thought, with additional axes being identified, and iii) latent spaces trained in a self-supervised manner are superior to those trained with supervised models, and generalize to other cell types and drug conditions without explicit training on those cell types or drug conditions. Future applications of MitoSpace include creating mitochondrial biomarkers for drug discovery and determining the effects of unknown drugs and diseases for diagnostic purposes.

## INTRODUCTION

The goal of high-content phenotypic screening^1,2^ is to use the link between the morphology of a cell and its state to identify drug candidates that move the cell’s state in a desired manner. The morphology of a cell can thus guide the identification of a disease and its underlying aberrant states and the identification of drugs to correct these states.

In traditional high-content phenotypic profiling^2,3^, a set of features is manually defined and quantified from each microscopic image. Examples include size, shape, and texture of an organelle^2,3,4,5^. Such feature sets can then be processed to find a subset of features that constitute a ‘fingerprint’ of each individual sample condition, for example healthy versus diseased. Prominent successful examples of this profiling technique include the NCI-60 tumor cell line panel^6^ where 60 different tumor cell lines were used to extract patterns of anticancer drug mechanisms^7^ and the relation between small molecule drugs, genes, and diseases^8^. Further examples include the determination of mechanism of action of pharmacological compounds^9, 10, 11,12,13,14^, target identification^15,16^, gene relationship identification^16,17,18,19,20,21,22^ and characterization of cellular heterogeneity^23,24^. Classic modeling techniques in this method, e.g. hierarchical clustering^25^, have in recent years been replaced with sophisticated deep learning methods in phenotypic drug development^26^. Specifically, convolutional neural networks (CNNs), have demonstrated superior classification performance through enhanced extraction of image-intrinsic features^27,28^. However, at the core of all of these approaches remains human supervision. Human annotation of training data and creation and curation of feature lists are used to train supervised machine learning algorithms (See Figure S1 left).

Advances in *self-supervised* deep learning have recently allowed the modeling of complex data distributions without explicit human input or the a-priori selection of features, for example for natural language processing^29,30^ and images^31,32,33^. Such methods have been applied before in the medical domain^34^, for example for protein structure prediction^35^, medical imaging^36^, and microscopy^37^. In a recent biological study, self-supervised deep learning was successfully applied to profile subcellular protein localization in microscopy data using autoencoders and a proxy classification task involving protein identifier labels^38^.

In this article, we present a new approach to phenotypic screening that is based on self-supervised deep learning. Specifically, our approach is not based on traditional cell profiling techniques but is rather based on novel ‘cell perceiving’ techniques that do not require the pre-definition of features nor the labeling of the data. i.e. there is no human or other input telling the model which image is associated with which drug treatment. The model learns the inherent similarity and dissimilarity between images using relative associations between data as opposed to absolute given labels. Also note that in cell perceiving, the self-supervised neural networks do not require feature selection nor feature extraction steps. The result is that human bias in feature conception, feature selection, and data annotation is eliminated. In our approach, the models detect these features themselves internally. We can thus map the true latent factors of variation in the data without human intervention.

The mitochondrial network stands out among the different organelles of a cell as one of the most diverse and responsive to cellular state and cellular health. Mitochondria are small organelles that provide up to 90% of a cell’s energy^39,40^ and are implicated in a vast array of diseases such as metabolic disorders^41^, developmental disabilities^42^, epilepsy^43^, neurodegenerative disease^44,45,46^, cancer^39,47^, and aging^45,47,48^. For that reason, mitochondrial morphology has been the target of high content phenotypic screening in cancer^49^, neurodevelopment^50^, metabolic diseases^51,52^, and neurodegeneration^53,54,55^ (comprehensively reviewed here^56^). These high-content screening studies are all of the ‘profiling’ type and follow the pattern that 1) hit parameters in the screen are defined, e.g. increase mitochondrial length by > 2.5 z-score^50^, 2) control cells and disease cells are exposed to a drug compound library, 3) imaged using fluorescence microscopy, and 4) the resulting images analyzed using software to quantify the hit desired parameters.

Given that the mitochondrial morphology is exceedingly complex, we hypothesized that manual definition of features and hit parameters was masking a multitude of factors that influence mitochondrial phenotype in response to drug candidates. Thus, important features would have remained so far unreported and unquantified.

In this study, we use contrastive and bootstrapping-based self-supervised deep learning approaches to create image-based semantically meaningful latent spaces which can act as a map for understanding mitochondrial phenotypes and for modeling the effect of pharmacological interventions. Our method requires no human labeling or feature definition. By not requiring human-defined classification labels, our approach is robust against overfitting and can correctly delineate generalizable factors of variation present in the data.

By focusing on drug-induced mitochondrial morphology and function changes we create an interpretable atlas of the accessible function/morphology space of the cell that links to cellular state and thereby drug function.

We show how the traditional dimension of mitochondrial phenotypic change, i.e. the change from fragmented to tubular, is augmented by additional dimensions that have so far remained hidden in previous manual or supervised approaches. We show how MitoSpace (i) captures meaningful functional and morphological information with no human supervision, (ii) generalizes across cell types and to drugs unseen during training, and (iii) is semantically superior to supervised methods. MitoSpace provides a powerful method to characterize and contextualize the effects of diseases, novel drugs, and other interventions on mitochondria.

## RESULTS

### Generating an Image Dataset that Captures the Full Variability of Drug-Induced Phenotypic Responses of Mitochondria

Mitochondria form a three dimensional network in the cell that changes over time through fission, fusion, and motility^5,39^. Consequently, it is established that the mitochondrial network naturally displays a wide range of morphologies in control conditions (**Figure 1A**). Beyond this cell to cell variability, it is established that mitochondrial morphology responds to stress or a metabolic switch to glycolysis with a punctate / fragmented morphology (**Figure 1B, left**) and to stress relief and a metabolic switch to oxidative phosphorylation with an elongated or hyperfused morphology (**Figure 1B, right**)^56,65^. However, there is a wide variety of additional reported mitochondrial morphology phenotypes^57^ that cannot easily be placed on the punctate-to-networked axis (see examples in **Figure 1C**). Morphologies like these have been widely observed but have traditionally been classified as outliers. We hypothesized that these mitochondrial morphologies belong to additional, currently uncharacterized dimensions of mitochondrial morphology responses that have so far remained hidden because of insufficient sampling of these states and manual analysis methodologies that predefined the goal observables to be along the punctate-to-networked axis.

**Figure 1.**
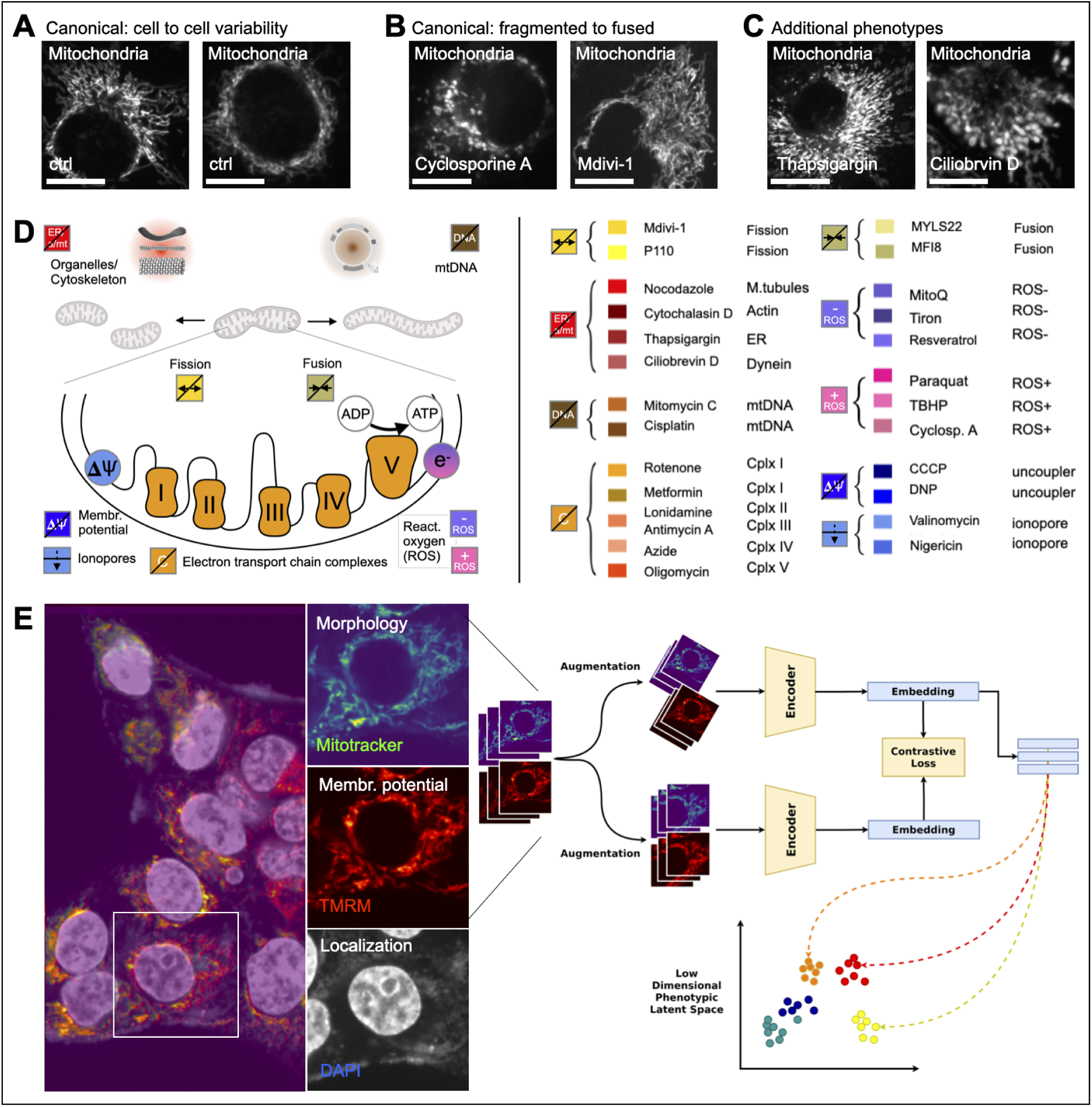
Self-supervised deep learning of drug-induced mitochondria phenotype changes with MitoSpace. (A) Mitochondrial network morphologies (white) are known to show a large cell to cell variability in control cells (HeLa) in control conditions (wt). (B) Mitochondrial networks (white) are reported to be distributed on an axis that ranges from highly fragmented (left) to highly fused (right). (C) Additional mitochondrial morphologies are hard to place on the fragmented-fused axis and have historically been hard to contextualize. (D) Depicted are the 9 major drug classes affecting mitochondria: i,ii) fission/fusion (yellow, beige), iii) interaction with other organelles such as the ER and the cytoskeleton (red), iv) mitochondrial DNA (brown), v,vi) mitochondrial membrane potential through uncouplers and ionophores (dark and light blue), vii) mitochondrial electron transport chain complexes (orange), and viii,ix) reduction (purple) or elevation (pink) of reactive oxygen species (ROS). 25 drugs belonging to these 9 drug classes were applied to cells to test the hypothesis that there are additional dimensions beyond the canonical fragmented to fused dimension. (E) High content fluorescence microscopy data of cells (HeLa, Cal27) was collected to capture mitochondrial morphology (Mitotracker, green), mitochondrial membrane potential (TMRM, red), and the location of the nucleus (DAPI, white) and were used to train a self-supervised deep learning model. After a data augmentation step, the model consists of two encoders that learn representations that aim to maximize agreement, thereby extracting signals that are common to entire classes of images while ignoring spurious factors such as background. The result is a reduced parameter representation of the cell images - an atlas that maps mitochondrial morphology to drug action.

To test our hypothesis, we set out to create an imaging dataset that contains the full variability of drug-induced mitochondria phenotype responses in order to contextualize them using self-supervised deep learning.

We used two cell lines, Cal27 head and neck squamous carcinoma cells^58^ and HeLa cervical cancer cells^59^, as model systems. Eight classes of mitochondria drugs were employed to induce the spectrum of mitochondrial morphology and function responses (**Figure 1D**): 1) fission inhibitors (Mdivi-1, P110); 2) fusion inhibitors (MYLS22, MFI8); 3) agents affecting mitochondrial motility (Nocodazole for microtubules, Cytochalastin D for actin filaments, and Ciliobrevin D for dynein) and other cellular organelles that interact with mitochondria (Thapsigargin for ER); 4) agents that affect mitochondrial DNA (Mitomycin C and Cisplatin); 5) agents that dissipate the mitochondrial membrane potential (CCCP and DNP); 6) agents that act as ionophores (Valinomycin and Nigericin); 7) agents that affect the mitochondrial electron transport chain complexes (Rotenone for Complex I, Lonidamine for Complex II, Antimycin A for Complex III, Azide for Complex IV and Oligomycin for Complex V); and agents that affect the production of reactive oxygen species (ROS) (Paraquat, TBHP, and Cyclosporine A to 8) induce ROS, MitoQ, Tiron and Resveratrol to 9) buffer ROS).

We used an automated confocal microscope to fluorescently image the mitochondrial morphology using Mitotracker Green and the mitochondrial function using the mitochondrial membrane potential marker TMRM. To facilitate automated cell segmentation we labeled cell nuclei using DAPI (see **Figure 1E**). For each cell line, we imaged ∼2,000 individual cells for each drug condition, resulting in a total of 2 cell lines x 25 drugs x 2,000 cells = 100,000 individual cell images. We used this dataset to generate our self-supervised deep learning-based mitochondria phenotype space.

### Self-supervised Deep Learning to Generate the Latent Space of Drug-Induced Mitochondrial Phenotypes

We use self-supervised contrastive learning to generate the latent space of drug-induced mitochondrial phenotypes, without requiring any human labeling. The ‘latent space’ in this context refers to a lower dimensional vector space (for example, lower than the 256×256 = 65,536 dimensional space of the images) that captures only the important factors of variation in the data.

We compared two self-supervised deep learning architectures, SimCLR^31^, a contrastive learning approach, and BYOL^32^, a related method leveraging bootstrapping, to learn semantically meaningful latent representations from our image dataset while ignoring spurious factors (see **Figure S1** for model architectures). Briefly, 256×256 images of single cells of both mitochondrial morphology (MitoTracker) and mitochondrial membrane potential (TMRM) were passed through a data augmentation step. We designed our data augmentations to explicitly make the model robust to potential spurious imaging and experimental factors such as noise and cell shape (see Methods Section). The data augmentation helps the network learn invariance to any spurious features which are not related to morphology/function, for example rotation of mitochondria, cell shape, translation, imaging noise, or brightness variations in the images. In the second critical step, the augmented images enter two separate encoder pipelines that are compared to each other so that contrastive loss between the images is minimized. The two heads share weights and are encouraged to learn consistent representations of the same cell image (and hence the same phenotype), irrespective of changes to the spurious factors mentioned above. The resulting embedding is the low dimensional phenotypic feature space that encodes and contextualizes mitochondrial morphology and function. As a result of this training, our model, MitoSpace, naturally learns the inherent phenotypic similarities and differences between different cells, grouping similar phenotypes together (**Figure 1E**). The resulting lower dimensional latent space is an atlas of drug-induced mitochondrial phenotypes which can act as a map for understanding and relating mitochondrial morphology to drug function.

### MitoSpace Delineates Phenotypic Activity from Images

The phenotypic atlas learnt by MitoSpace for Cal27 cells is shown in **Figure 2**. Our method is learning phenotypic similarity, i.e. locating similar phenotypes close together in space. Being at one point in the map and moving gradually into one direction we would expect the mitochondrial morphology to gradually change and morph into a new phenotype. We used UMAP^59^ to reduce the 128 dimensional latent space learnt by the model down to 3 dimensions for visualization purposes. Note that every conclusion in the paper is directly drawn from the 128 dimensional space and that UMAP is used for easier visualization only. Each point in this latent space represents an image of a single control or drug-treated cell. The position of the point and its relation to all the other points (images) reflects the model’s understanding of the morphological (MitoTracker) and functional (TMRM) phenotypes expressed in that cell image in relation to the other points. Each point is colored by the drug that the cell was treated with. Colors are chosen for each drug based on the drug’s reported mechanism of action (**Figure 2A**). We found that MitoSpace reliably maps the semantic landscape of drug-induced latent mitochondrial phenotypes, grouping drugs with similar effects on mitochondria together. This can be observed based on the color coding in the latent space, where drugs that have similar mechanisms of action are colored by a similar color (see also **Figure S2** for the latent spaces of different methods).

**Figure 2.**
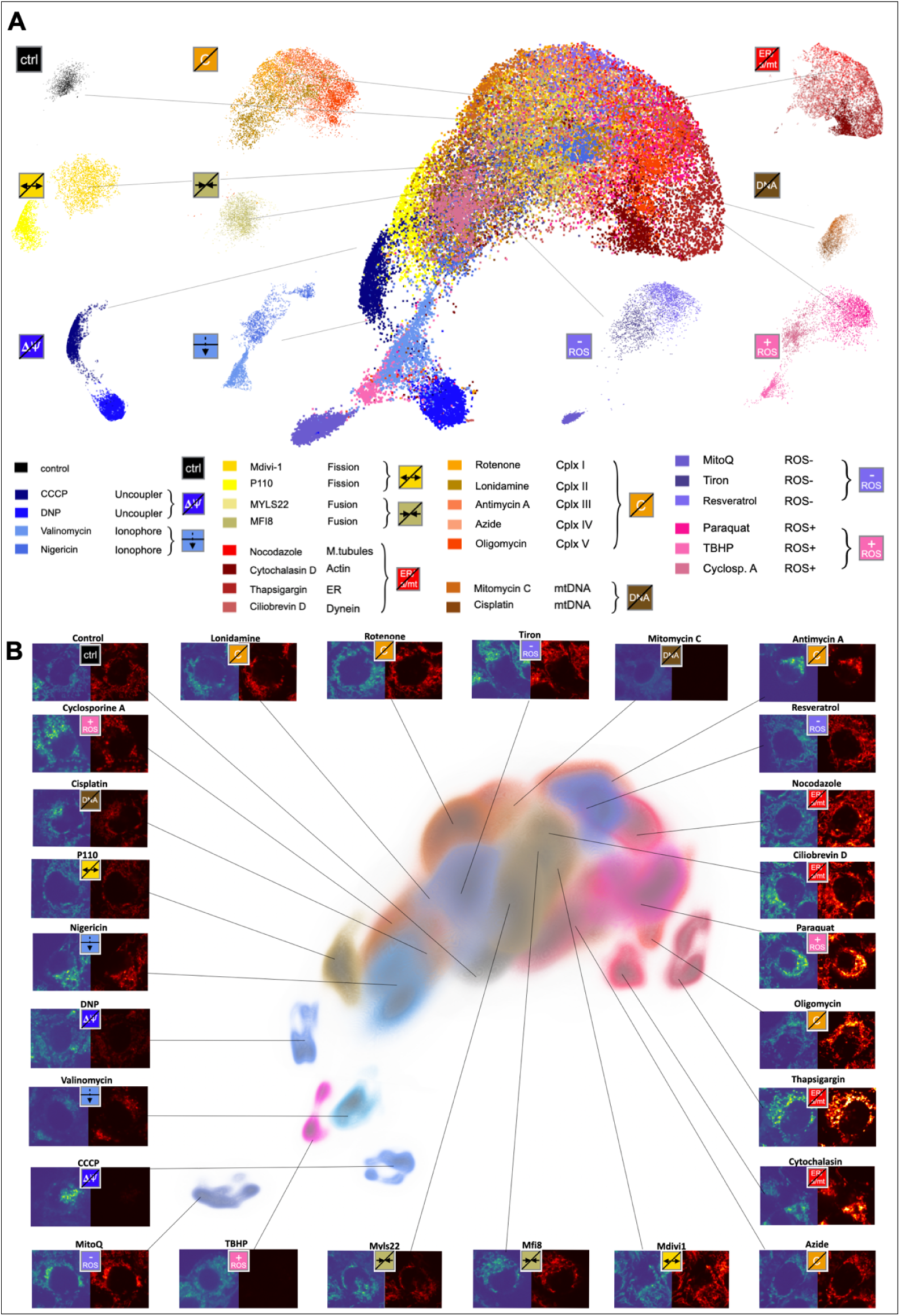
MitoSpace creates the first atlas of drug-induced mitochondrial phenotypes. (A) Depicted is our image-extracted semantically meaningful latent space of drug-induced mitochondrial phenotypes. Every pixel represents a cell image taken under control or drug treatment conditions. A larger distance between images indicates greater differences in mitochondrial phenotype. Note how drugs of the same class (e.g. uncouplers, ionophores, agents that affect the respiratory chain complexes, etc.) cluster in the same regions of the space, indicating that mitochondria react to similar treatment similarly and that MitoSpace can semantically extract these similarities. (B) Representative images of mitochondrial network phenotypes for each cluster are shown in this panel (green, mitochondrial mass, mitotracker; red, mitochondrial membrane potential, TMRM). Note how similar morphologies (e.g. punctate for uncouplers and ionophores in the bottom left) and similar membrane potentials (e.g. high membrane potentials in hyperpolarized, oligomycin-treated cells on the right side of the space) cluster together in the space.

We then analyzed the latent phenotypes more closely and found that the atlas self-organizes to capture a wide variety of phenotypes inherent in the data. The different drug conditions are found to be placed radiating into different directions from a common center in the space where the control condition is found. This indicates that these drugs have resulted in quantifiably different mitochondrial phenotypes that are distinct from each other. **Figure 2B** depicts MitoSpace in the form of a density plot to aid visualization surrounded by representative cell images taken from the center of each drug cluster. The center or mode of each cluster is where the most characteristic images from each drug condition are located. We found that each image cluster shows clear differences in mitochondrial morphology. The control condition images are found clustered at the center of the latent space and show the baseline mitochondrial morphology and membrane potential. In drug conditions that affect the mitochondrial membrane potential through uncoupling (CCCP, DNP) and through acting as ionophores (Valinomycin, Nigericin), mitochondria show punctate morphology and low membrane potential (**Fig. 2B left**). Fission (Mdivi-1, P110) and fusion inhibitors (MYLS22, MFI8) cluster close to the control condition in the center of the space and show more and less punctate morphology, respectively, and close to control membrane potential (**Fig. 2B bottom**). Images taken under mitochondrial motility affecting drugs (Nocodazole, Cytochalastin D, Ciliobrevin D) and other organelle affecting drugs (Thapsigargin) cluster to the right side of the space, showing a more filamentous mitochondrial structure and high mitochondrial membrane potential (**Fig. 2B right**). Images taken under the effect of agents that affect the electron transport chain complexes (Rotenone, Lonidamine, Antimycin A, Azide, Oligomycin) cluster to the top right of the space (**Fig. 2B top right**), and images taken under mitochondrial DNA affecting drugs cluster to the top left of the space (**Fig. 2B top left**). Agents that affect the production of reactive oxygen species (Paraquat, TBHP, and Cyclosporine A, MitoQ, Tiron, Resveratrol) were found to be clustered in both ends of the space. Overall, we could observe that our model clustered similar morphologies and similar mitochondrial function conditions in similar regions of the space. This means that cells are mapped in the direction of the respective phenotypes in varying degrees, correlating to how strongly they are affected by a drug intervention.

Not being constrained by human labels, we find that MitoSpace focuses on actual phenotypic responses of cells. This means that MitoSpace does not overfit to try and conform to predefined labels. This is important because even cells treated with the same drug may show a range of phenotypes, with some cells responding mildly and other cells responding strongly to the same drug. Not enforcing the same label on such widely responding cells allowed us to capture the true response of each condition, resulting in a multidimensional continuum of phenotypes where each cell image is mapped to its correct phenotypic position irrespective of what drug it is treated with. In a later section, we show how supervised models trained on the same data, constrained by predefined labels, suffer from overfitting and are hence unable to create semantically meaningful latent spaces. Such automatic feature selection, extraction, and mapping are the keystones of the ‘cell perceiving’ approach as compared to ‘cell profiling’.

### MitoSpace Introduces Principal Semantic Directions that Combine Form and Function of Mitochondria

We now show how MitoSpace captures various phenotypic features inherent in the data, without having to manually define them. We hypothesized that there would be mitochondrial axes beyond the established punctate-to-networked axis, and self-supervision would allow us to capture these. Note that, while we create an association between known mitochondrial phenotypic features and the latent space, it is important to recognize that the deep learning model may be capturing higher order features that may not be easily defined by simple features previously defined in the literature. This analysis is thus meant to be representative, not exhaustive.

We conducted principal component analysis (PCA) in the latent space generated by the MitoSpace model in order to discover phenotypic axes and to gain deeper insights into the connections and clusters that MitoSpace found in mitochondrial phenotype space (**Figure 3**). We found that > 70% of our variance in the latent space was accounted for by the first 10 principal components (PCs) and > 55% by the first 5 PCs (**Figure 3A**). We computed the correlation of these PCs with independently computed mitochondrial features extracted from each cell image. Specifically, we measured correlation with (i) the fluorescence intensity of MitoTracker and TMRM, and (ii) ten morphology and network features extracted using Mitochondria Analyzer^60^ (**Figure 3B**). Specifically, we quantified total branch length per mitochondria, mean form factor, number of branch junctions, number of branch end-points per mitochondrion, total area of the mitochondrial network per cell, total mitochondrial branch length, number of mitochondrial network branches, total perimeter of the mitochondrial network, number of mitochondria per cell, and mean aspect ratio of mitochondria. **Figure 3C** shows the Pearson correlation coefficient (PCC) of these independently extracted features with the PCs extracted from the learned latent space (see also **Figure S4** for PCCs of the other methods). We find that MitoSpace captures the variation in these features, in spite of there being no explicit feature selection step anywhere in our pipeline. This is reflected in the high and statistically significant correlation coefficients in **Figure 3C**. For the first PC, three features can be highlighted (**Figure 3C, star**) that are closest to the reported axes along which mitochondrial phenotype and function are arranged: a) TMRM fluorescence intensity which has been reported to correlate with mitochondrial membrane potential, b) mitochondrial aspect ratio, a feature that closely tracks the punctate-to-networked axes of mitochondrial morphology, and c) mitochondrial area. For the first PC, we observed that TMRM is not correlated, aspect ratio is somewhat negatively correlated, and mitochondrial area is strongly negatively correlated. This pattern however is different for each PC and for some, e.g. PC-3, it is reversed with TMRM being strongly positively correlated and aspect ratio exhibiting a correlation close to zero. We hypothesized that while the individual PCs might show one or the other pattern or correlation, the full MitoSpace model which integrates the entire variance of phenotypes available in our dataset would exhibit recognizable patterns. We plotted the intensity of each feature for each image as a color gradient on top of the MitoSpace-generated latent space and found that such patterns are indeed recognizable (**Figure 3D**). TMRM shows a clear trend from low TMRM intensity at the bottom left end of the latent space to high intensity at the right end of the latent space (**Figure 3D, left**). A similarly clear patterning exists for mitochondrial area from low mitochondrial area (top) to high mitochondrial area (bottom) of the latent space (**Figure 3D, middle**), and for mitochondrial aspect ratio (low aspect ratio in the bottom left to high aspect ratio in the top right, **3D, right**). These continuous feature variations in the latent space can be thought of as novel axes that encode simultaneous descriptions of mitochondrial form and function that augment traditional ‘punctate-to-networked’ axes into a much richer description of mitochondrial phenotypes. To probe these axes we sampled along their gradients to show representative images (**Figure E**). Indeed, the TMRM axis embedded in the latent space shows not only the feature itself (continuously decreasing TMRM intensity, red signal, **3E, bottom**) but simultaneously shows a mitochondria morphology continuum from more punctate to networked morphologies. In contrast, sampling along the mitochondrial area axis shows relatively constant TMRM intensity and network morphology while it is mainly the mitochondrial area that changes (**Fig. 3F**). Finally, sampling along the mitochondrial aspect ratio feature axis in the latent space shows the expected reticular to punctate morphology gradient while TMRM signal remains constant (**Fig. 3G**).

**Figure 3.**
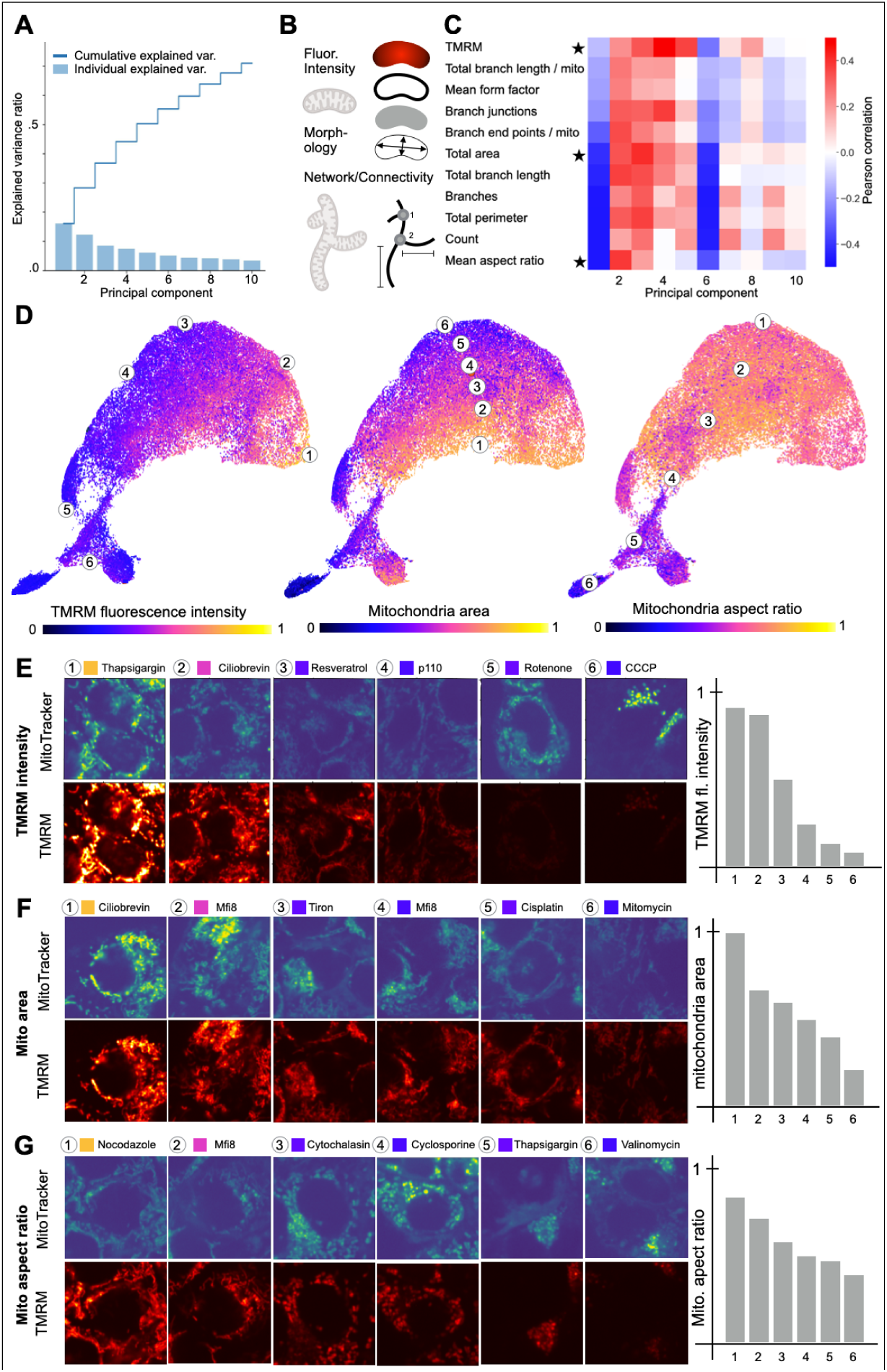
MitoSpace introduces semantic directions that integrate both mitochondrial form and function. (A) Principal component (PC) analysis of MitoSpace shows that the first five PCs (blue bars) explain 58% of the variance in the dataset (light blue line, cumulative variance). (B) To interpret MitoSpace’s PCs, we quantified the mitochondrial network with respect to i) fluorescence intensity of MitoTracker and TMRM, and morphometric features of ii) mitochondrial network fragments and iii) the mitochondrial networks. (C) We found that different principal components (PCs) correlate differently with different features that together show the emergence of a pattern. Traditional semantic directions (axes) such as TMRM intensity axes or punctate-to-networked axes (aspect ratio) are highlighted with a star. (D) We find that traditional axes of mitochondrial characterization such as i) mitochondrial membrane potential (TMRM), ii) mitochondrial area, and iii) mitochondrial aspect ratio are preserved in MitoSpace. Clusters of morphological responses to individual drugs appear as superpositions of multiple axes. Note the clear patterning in form of a gradient that is present for each depicted mitochondrial feature. MitoSpace integrates all of them into a unified semantically meaningful space. (E) Mitochondria membrane potential axis: Note how the TMRM signal along this axis changes from very high intensity to medium and eventually to low intensity, indicating that the mitochondrial membrane potential is vanishing along this axis. Also note how the mitochondrial morphology changes from highly tubular networks to small punctate structures where the mitochondrial membrane potential is lowest. (F) Mitochondria area axis: Note how the mitochondrial area in the MitoTracker channel changes from large volumes of the cytoplasm covered by mitochondrial density to medium and then low volume of mitochondrial density. Also note that this change is not accompanied by a drastic change in TMRM intensity indicating that the mitochondrial membrane potential is less important in this axis. (G) Mitochondria aspect ratio axis: Note how the mitochondrial morphology changes from a tubular reticulon, to a mixture of tubules and fragments, to a very fragmented network along this axis.

Together, these results show how MitoSpace directly captures and quantifies human-interpretable and previously validated features related to mitochondrial morphology and function, which lends validity to its semantic structure. Within this semantic structure, each drug class has its own location where the combination of mitochondrial morphology features and function (TMRM) act as a unique identifier that would allow the characterization of an acting drug from images alone.

### MitoSpace Accurately Captures the Semantic Meaning of Drug-Induced Mitochondria Phenotypes

Next we were interested in the accuracy of MitoSpace in predicting drugs from images alone. We note that since each cell may react differently even to the same pharmacological intervention, drug labels and drug function labels may not always be truly representative of the phenotype expressed in each cell image. Each cell may express multiple phenotypes in various proportions in spite of the drug it is treated with, and hence the true latent space is more akin to a mixture model over latent phenotypes rather than points with hard drug label assignments. Self-supervised models allow us to capture this latent phenotype of each cell, independent of which drug it is treated with. This is a key factor that allows the construction of biologically and semantically meaningful latent spaces. However, it also makes the models hard to evaluate. Hard classification accuracy based on drug labels as a metric does not fully capture the semantic validity of the generated latent space. Cognizant of this, we used top-3 accuracy as a proxy metric for evaluating performance of the latent space taking into account its mixed nature. We obtained above 84% top-3 drug-ID accuracy and close to 86% drug-class accuracy with our chosen model BYOL (see **Table 1, row 1, Figure S5**). This indicates that the model correctly places each cell image in the latent space close to the mode of the pharmacological intervention it is treated with. We tested if other self-supervised model architectures would yield higher accuracies for MitoSpace but that was not the case. Slightly lower accuracies were obtained with SimCLR, with a more recent transformer-based approach, DINO^33^, and with an autoencoder-based approach (a VQVAE with a classifier head^38^) (**Table 1, rows 2-4**).

**Table 1.** MitoSpace is highly accurate at predicting drugs from images while simultaneously creating a semantically meaningful latent space. Displayed are top 1 and top 3 prediction accuracies for individual drugs, top 1 and top 3 accuracies for drug function, and our semantic quality metric for MitoSpace and other model architectures. Note how supervised models achieve slightly higher accuracies in raw prediction tasks but fail at the task of transfer learning, i.e. creating a semantically/biologically meaningful latent space that is interpretable (high semantic quality scores). Also note that the supervised KNN model on mitochondria analyzer features does not have a latent space and thus a ‘semantic quality metric’ cannot be computed.

| Model | Top 1 Drug Accuracy | Top 3 Drug Accuracy | Top 1 Drug Function Accuracy | Top 3 Drug Function Accuracy | Semantic Quality Metric |
| --- | --- | --- | --- | --- | --- |
| *Self-Supervised* |  |  |  |  |  |
| **MitoSpace BYOL** | **0.609** | **0.841** | **0.645** | **0.859** | **0.217** |
| MitoSpace SimCLR | 0.601 | 0.814 | 0.639 | 0.856 | 0.207 |
| MitoSpace VQVAE w/ proxy classifier | 0.412 | 0.703 | 0.469 | 0.736 | 0.173 |
| MitoSpace DINO | 0.291 | 0.574 | 0.359 | 0.620 | 0.156 |
| *Supervised* |  |  |  |  |  |
| Baseline Resnet18 | 0.730 | 0.916 | 0.745 | 0.926 | 0.153 |
| Mitochondria Analyzer | 0.057 | 0.180 | 0.135 | 0.240 | - |

Next, we wanted to compare our self-supervised approach to traditional, supervised approaches. As a comparison, we trained a baseline supervised ResNet18^61^ on our dataset and also included a naive K-nearest neighbors (KNN) model trained on a set of 17 features previously defined in literature, extracted from each image using Mitochondria Analyzer^60^. While the KNN performs expectedly poorly (18% and 24% drug-ID and drug-class performance respectively), the ResNet18 model performs very well and better than our self-supervised approaches (92% and 93% drug-ID and drug-class performance respectively) (**Table 1, rows 5-6**).

This is not surprising. Self-supervised models typically show slightly lower classification performance than supervised models on standard image benchmarks like ImageNet-1K. However, they are known to exhibit better transfer learning performance^66,67^, that is, they generalize better to concepts and data that are out-of-distribution which the model has not seen at training time. This is especially important for a phenotypic atlas like MitoSpace, where the effects of unseen drugs and the resulting phenotypes must be accurately captured.

To show the superior transfer learning performance of MitoSpace compared to ResNet18 and its superior performance as a phenotypic atlas, we plotted the test-set latent spaces generated by MitoSpace and ResNet18 (**Figure 4**). MitoSpace’s latent space shows a continuous map in which individual images of drug treated cells blend into each other (**Fig 4A**). In contrast, the ResNet18 latent space shows each drug cluster separated from each other with large distances in between them (**Fig 4B**). This structure reflects the supervised model’s superior performance in raw accuracy, but it comes at the expense of lost semantic/biological meaning. We demonstrate two situations to make this clearer.

**Figure 4.**
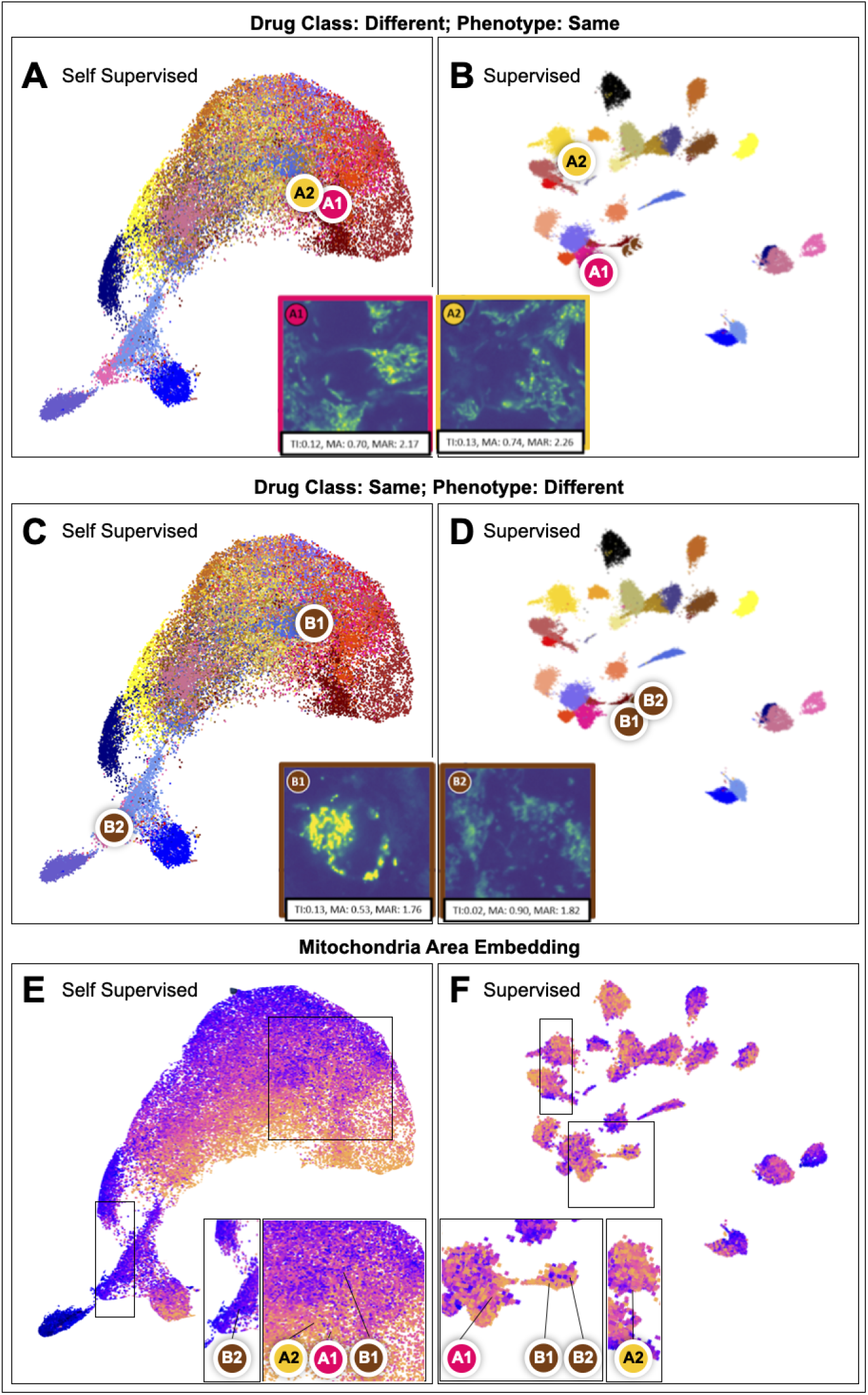
Self-supervised models generate higher quality latent spaces than supervised models. (A) Self-supervised models correctly place cells A1 and A2, with similar phenotype, together, in spite of them being treated with different drugs. (B) Supervised models overfit to predefined drug labels and incorrectly place A1 and A2 away from each other even though they have the same phenotype. (C) Self-supervised models correctly separate cells B1 and B2, even though they belong to the same drug, since they have different phenotypes, and place them close to other cells with similar respective phenotypes. (D) Supervised models place B1 and B2 close together, i.e. they pack all cells treated with the same drug together even if they respond differently (E) Variation of the Mitochondria Area over the self-supervised latent space. Note how there is a clear semantic pattern (color gradient). (F) Variation of the Mitochondria Area over the supervised latent space. Note how there is no clear semantic pattern but that rather each cluster groups drastically different phenotypes together.

First, consider two cells, A1 and A2, treated with different drugs, but having the same phenotypic values (A1: TMRM intensity=0.12, Mean Mitochondria Area=0.70, Mean Mitochondria Aspect Ratio=2.17, A2: TMRM intensity=0.13, Mean Mitochondria Area=0.74, Mean Mitochondria Aspect Ratio=2.26). Since the self-supervised model placed cells close to any other cells showing similar phenotype irrespective of what drug they are treated with, it correctly mapped A1 and A2 close together (**Figure 4A**). The supervised model, on the other hand, overfitted on the predefined drug label and incorrectly put A1 and A2 far away from each other (**Figure 4B**). Second, consider two cells, B1 and B2, treated with the same drug but showing completely different phenotypes (B1: TMRM intensity=0.13, Mean Mitochondria Area=0.53, Mean Mitochondria Aspect Ratio=1.76, B2: TMRM intensity=0.02, Mean Mitochondria Area=0.90, Mean Mitochondria Aspect Ratio=1.82). The self-supervised model, again correctly placed B1 and B2 far away from each other, closer to other cells showing phenotypes similar to each of them (**Figure 4C**). The supervised model again overfitted on the predefined drug label and incorrectly placed these images close together (**Figure 4C**). While we only showcased two examples, such behavior is prevalent in all our supervised and self-supervised models.

While it is now apparent that the supervised latent space is not meaningful in terms of pharmacological function (i.e. it does not map drugs with similar function close together), it does not correctly capture individual features either.

**Figure 4E** shows the variation of a particular feature, the mean mitochondria area, over MitoSpace’s self-supervised latent space. We can observe how the consistent variation of the feature is correctly captured by the model as discussed previously. In contrast, the same feature is plotted on the supervised latent space in **Figure 4F**. We can see that there is no consistent gradient over the latent space that captures the variation of the feature. Instead, cells with different mitochondrial areas are mixed together without any particular pattern. The model gives priority to the drug label provided during supervision as opposed to true underlying phenotype. Essentially, supervised classification models overfit to predefined drug labels leading to a latent space with inferior semantic/biological meaning. Thus, they may not generalize to out-of-distribution drugs or cell types, and no qualitative or quantitative biological conclusions can be drawn from latent spaces generated in such a manner.

To quantitatively capture this semantic quality of the latent spaces of self-supervised models, we computed a semantic quality metric for each model. The semantic quality metric quantifies how well the latent space captures previously known features related to mitochondria. We define this as the aggregate Pearson Correlation Coefficient of the first 10 principal components of the latent space with each of the 17 features extracted from Mitochondria Analyzer. Intuitively, the higher the semantic quality metric, the better the latent space captures these predefined features, i.e. its directions correlate better with semantic biological concepts. The metric is provided for the different models in **Table 1**. All self-supervised models showed higher semantic quality as compared to the supervised model.

### MitoSpace Generalizes to Different Cell Types and Drug Interventions without Explicit Training

Next, we investigated the generalizability of MitoSpace in two ways: (a) generalization to a different but similar cell type, and (b) generalization to drugs unseen during model training. Mitochondria can be found in all eukaryotic cells and, depending on cell type, assume different or similar roles between different cell types. On that basis, we hypothesized that our model would generalize across different cell types to the extent of the similarity between the cell types. We chose to investigate generalizability of trained models between Cal27 and HeLa cells. Similar to Cal27, HeLa is an immortalized cancer cell line that heavily relies on oxidative phosphorylation in mitochondria for their large energy demand. Different from the head and neck cancer cell line Cal27 however, HeLa has been derived from cervical cancer.

To investigate generalization across cell types, we trained models on one cell type and predicted all images of the alternate cell type using that model, i.e. we trained models using HeLa cells and predicted Cal27 cells and vice-versa. We compare the top-3 accuracy of the base model (the model trained on the same cell type) to the alternate model (the model trained on the other cell type). We also compared the resultant latent spaces visually.

**Figure 5** shows the results of our generalization experiments based on the drug-induced imaging dataset for Cal27 cells (**Figure 5A)** and HeLa cells (**Figure 5B**). **Figure 5C** and **Figure 5D** show the predicted Cal27 latent space for all drugs in the form of density plots, for models trained on Cal27 cell images (i.e. the base model) and HeLa cell images (i.e. the alternate model) respectively. While we did find minor differences, we found that the overall structure of the space is maintained, both globally and locally, irrespective of which cell type the model is trained on. Note that the deep learning latent spaces are quite robust and reproducible (see the consistent latent spaces in **Figs 5 C, D, E, F,** in addition to the latent spaces for various different methods in Figures S2). The UMAP visualization can be reproduced by fixing the random seed. Note that the UMAP embedding is for visualization purposes only. Any downstream analytical conclusions are drawn directly from the 128 dimensional latent space.

**Figure 5.**
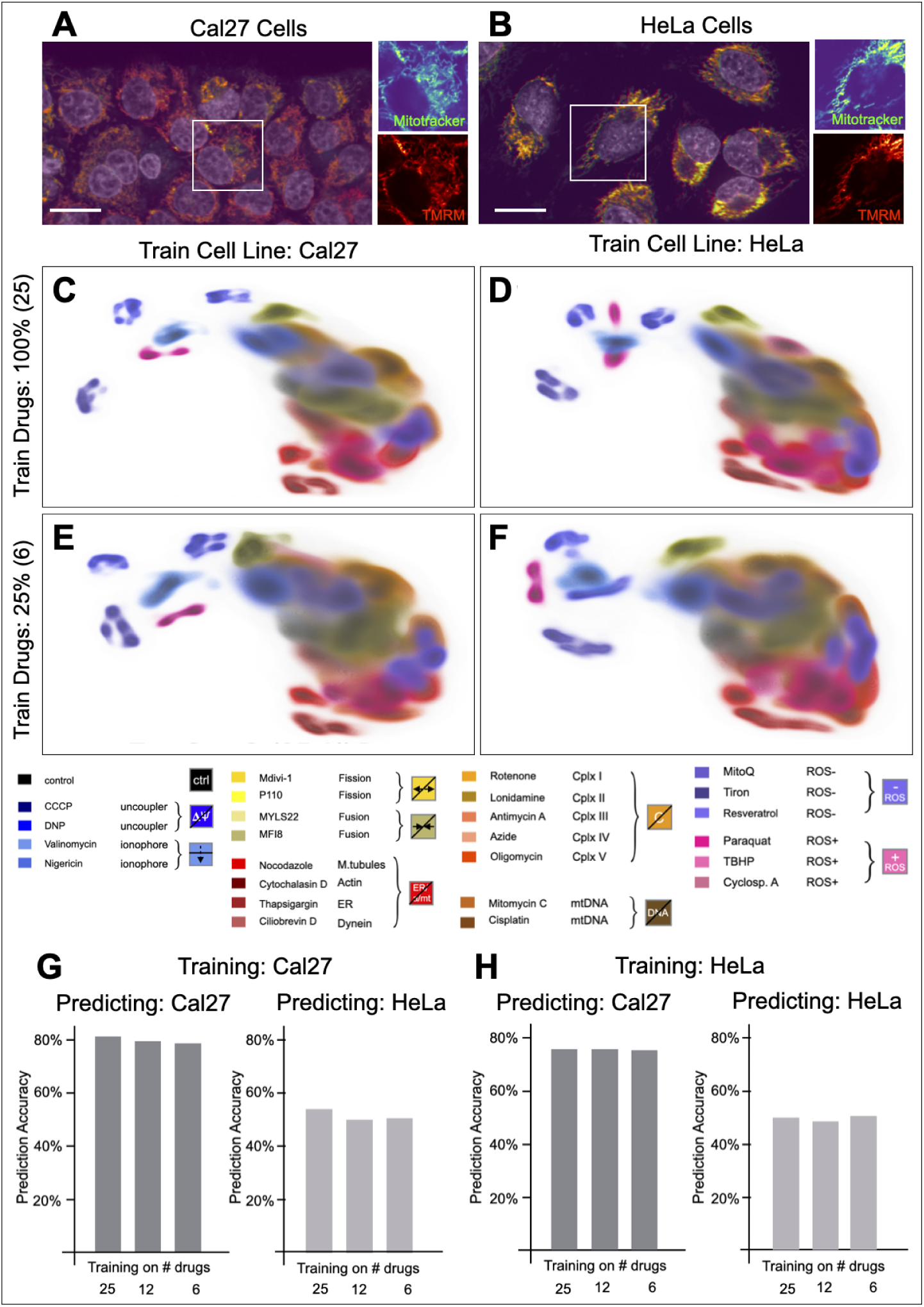
Mitospace generalizes to different cell types and drugs without explicit training. (A) Cal27 HNSCC cells form tightly packed colonies from compact polygonal cells. (B) HeLa cells form loose assemblies from individual spread-out polygonal cells. (C) Predicted Cal27 mitochondria latent space using a model trained on the same Cal27 cell images using 100% of the training data. (D) Predicted Cal27 mitochondria latent space using a model that has been trained on HeLa cell images using 100% of the training data. Note that the predicted latent spaces trained on Cal27 and HeLa are remarkably similar. (E) Predicted Cal27 mitochondria latent space trained on only 25% of Cal27 training data. I.e. predict 24 mitochondria drug responses in Cal27 cells using a space trained on 6 mitochondria drug responses in Cal27 cells. Note how similar the predicted locations are between C and E even though the model has only been trained on a quarter of the drug conditions. (F) Predicted Cal27 mitochondria latent space trained on only 25% of HeLa training data. I.e. predict 24 mitochondria drug responses in Cal27 cells using a space trained on 6 mitochondria drug responses in HeLa cells. Note the similarity between E and F and D and F, even though different cells were used for training and only a quarter of that data was used for training. (G) Prediction accuracy if the ground truth drug morphology is contained in the top three predictions given by the model (top-3) when the model is trained on Cal27 data. Cal27 predictions are consistently around ∼80% even if the number of drugs used for training is reduced. HeLa predictions are consistently around 57% even if the number of drugs used for training is reduced. (H) Top-3 Prediction accuracy of Cal27 and HeLa drugs based on the mitochondrial morphology in HeLa cells. A similar pattern is observed as in G. Note that the prediction accuracy falls only by ∼5% when training and prediction are not done on the same cell line.

To investigate generalization to drugs unseen during training, we withheld drugs during training, and predicted the entire set of drugs during testing. We then compared the resulting latent space to the oracle model that has access to all drugs during training. We train with as little as a quarter of our total drug library, ensuring that we choose our training drug set so that we still have adequate representation from each functional category.

**Figure 5E** and **Figure 5F** show the same Cal27 latent spaces trained on the two different cell types, but only using a quarter of the drugs at training time (but predicting all the drugs at test time). The reduced training set of drugs was chosen such that the set still maximally represented the extent of the latent space. Once again, the structure of the latent space was maintained. **Figure 5G** shows the top-3 accuracy of predicting each image into the correct drug category using a KNN predictor on top of the embeddings, for models trained on different cell types and number of drugs. We found only minor drops in accuracy when the underlying model is trained on a smaller number of drugs. As an example, prediction accuracy for train Cal27 test Cal27 drops from 81% to 78% when the number of training drugs is reduced from 24 to only 6 (**Figure 5G left**). Similarly, prediction accuracy for train Cal27 test HeLa drops from 56% to 51% when the number of training drugs is reduced (**Figure 5G right**). It is worth noting that our prediction success for Cal27 was high even when trained on HeLa data (**Fig. 5G,H, left**) while our prediction success for HeLa remained low even when trained on HeLa (**Fig. 5G,H right**).

In summary, our results indicate two advantages of our approach: (i) latent spaces trained using our method generalize across cell types, i.e. a model can be trained on a different but related cell type and it still shows meaningful results on the original cell type, and (ii) global and local structure of the latent space and quantitative measures like accuracy are maintained even when only a subset of drugs is used for training, implying that models can generalize to drugs unseen during training.

Such out-of-distribution generalization capability is possible because of the completely unsupervised nature of our cell perceiving method, and opens up possibilities for understanding behavior of a wide range of cell types with minimal data collection and deep learning model complexity. It would also allow the delineation of similarities and differences in pharmacological effects and therapeutic interventions across cell types.

## DISCUSSION

### A Novel ‘Cell Perceiving’ Approach Based on Self-Supervised Deep Learning to Map Mitochondrial Phenotype to Drug Effect

We have developed a novel approach using self-supervised deep learning to create a one-to-one mapping of mitochondrial form and function to drug effect, which we call MitoSpace. In addition, we introduce a new perspective, called ‘cell perceiving’, as opposed to traditional ‘cell profiling’. Cell perceiving is characterized by automated extraction and contextualization of phenotypic information from microscopic images without requiring human-defined labels or feature lists. We can thus create a phenotypic latent space without requiring any additional information beyond the images themselves. This latent space acts as a phenotypic atlas that reliably clusters cells treated with functionally similar drugs into the same areas and captures all mitochondrial phenotypic features present in the data without any explicit human-defined feature selection steps. This includes mitochondrial features previously defined in literature but may also include other, more complex features inherent in the data that cannot be easily characterized manually.

Limiting the set of features to those defined by human experts may not capture all dimensions of phenotypic variation present in the data. We demonstrate this by showing that a model based on human-defined features captured using Mitochondria Analyzer does not capture the semantic meaning of the combined mitochondria form and function space. In fact, biasing a model using manual drug labels may hide true phenotypes expressed by individual cells and may lead to overfitting.

### Latent Spaces Trained in a Self-Supervised Manner are Biologically Meaningful

We showed that phenotypic latent spaces obtained using self-supervised models are biologically meaningful. Specifically, we validated that (i) the latent spaces reliably capture and delineate previously defined features related to mitochondrial phenotype in a semantically meaningful way, with cells having similar feature values being mapped close together and vice versa (i.e. the latent space is locally meaningful), and (ii) cells treated with functionally similar drugs are mapped to the same area (i.e. the space is globally meaningful).

The overall location of drug-induced morphology clusters in MitoSpace makes sense in that regard where, for example, membrane uncouplers and ionophores are both mapped in a similar region of MitoSpace. Locations that are at first glance unexpected turn out to be supported by the literature. For example, MitoQ can cause both mitochondrial swelling and depolarization in some cell types^62^.

Such biological validity opens up avenues for defining computational biomarkers that can characterize all phenotypes expressed by a cell or organelle without manually defining them. For example, we can choose the principal components of the resulting latent space as the major directions of variation in the data, and represent each cell by the values of the first N principal components at that point.

### MitoSpace is Generalizable to Other Organelles, Cell Types, and Drug Libraries

Our method is independent of cell type, cell organelle, and drug library used and can be extended without any human annotation or effort except for sample preparation and image acquisition. We validate this by showing how the semantic structure and classification performance of our obtained latent space remains consistent when trained with a different cell type or when withholding drugs during the training phase in order to simulate unseen drugs during the test phase.

Given its generalizability, MitoSpace and similar latent spaces may be useful to determine the likely function of unknown drugs based on their effect on mitochondrial phenotype, as well as to identify the functional effect of diseases on mitochondria. They could be used for in-vitro pharmacology and as endpoints in clinical trials.

Beyond drug discovery and disease profiling, MitoSpace, and more broadly ‘cell perceiving’, offers exciting opportunities for basic research. The distinct mitochondrial morphologies that appear in response to specific drug effects likely result from unique signaling pathways that optimize cellular health under stress conditions. MitoSpace provides a window into these pathways and opens up avenues for further research.

### Practical Use of MitoSpace for Drug Discovery and Diagnostics

There are practical use cases for MitoSpace in drug characterization, drug discovery and diagnostics. 1) Drug characterization: Novel drug candidates for mitochondria typically have to undergo laborious and costly laboratory tests to determine if mitochondria are targeted by the drug candidate and which mitochondrial subsystem is targeted. With MitoSpace, microscopy images of cells under the influence of the novel drug can be taken and then placed into the latent space. The resulting location of the novel drug’s images in MitoSpace will identify the closest drug analogs as the closest neighbors in the latent space, thereby identifying the likely mechanism of action of the novel drug. 2) Drug discovery: For high-content library screening, a region of interest is defined in the latent space that is characterized by certain pharmacological properties (e.g. mitoprotection). In the next step, all images of drug candidates that are located within the region of interest are likely candidates for drugs that possess the desired properties. 3) Diagnostics: In the clinic, characterization of mitochondrial diseases is often difficult and requires multiple steps that involve blood draws and sequencing. With MitoSpace, microscopy images of a patient’s cells can be taken and placed into the latent space. The resulting latent space neighborhood will be images of disease phenotypes that are closest to the condition at hand, allowing a more focused screening and accelerated diagnosis.

### Limitations of our Approach

Limitations of self-supervised deep learning methods are in the requirement to carefully control the experimental conditions so that all potential human and technical error sources are either eliminated or well represented throughout the data. As a result, automated microscopy and automated sample preparation are likely required to maintain low noise levels while obtaining the large quantities of microscopy images required to train these models.

### Further Development of MitoSpace

Although powerful, multiple avenues exist to improve MitoSpace. Specifically extension to incorporate additional drugs, additional cell lines, additional organelles, and additional dimensions are the first directions of expansion.

Currently, our MitoSpace model uses 25 drugs as the basis for its applications. However, expanding the scope of drugs under study could significantly enhance the potential of our tool. By testing a broader range of drugs, we could potentially uncover a wider array of effects on organelle morphology and function, leading to novel insights into drug mechanisms and potential therapeutic targets.

Our work up to this point has utilized a limited selection of cell lines. To improve the robustness and generalizability of our findings, the next step is to incorporate a larger and more diverse set of cell lines into our methodology. Different cell types can react differently to the same drugs, making it crucial to understand these variable responses across cell lines.

While the application of MitoSpace to mitochondria provides a strong proof of concept, we hypothesize that other organelles could also benefit from this technology. Certain organelles like the endoplasmic reticulum and the cytoskeleton present unique complexities in their morphology that pose similar challenges for classical profiling methods as mitochondria. By extending MitoSpace to these organelles, we could provide more in-depth and accurate phenotypical quantification, helping researchers better understand their functions and interactions within the cell.

MitoSpace has so far been used to analyze two-dimensional images, but organelles are inherently three-dimensional structures that change over the fourth dimension of time. Therefore, integrating three-dimensional imaging and time-lapse movies into MitoSpace could significantly increase its predictive power. This expansion would enable us to capture the dynamic nature of organelles and better understand how they behave and interact under perturbations on cellular states.

## STAR METHODS

### Cell Culture

Cal27 and HeLa cells were cultured in 89% DMEM, high glucose, GlutaMAX (Gibco, #10566016), 10% FBS (GenClone, #25-550), 1% Penicillin-Streptomycin (Gibco, #15140122). Cells were washed with DPBS (Gibco, #14190144) and detached with 0.05% Trypsin-EDTA (Gibco, #25300054) before passaging onto imaging plates.

### Live Cell Imaging

Cells were passaged onto 96-well #1.5 glass bottom plates (Cellvis, #P96-1.5H-N) and grown for one day to reach a final density of 60-80%. After drug treatments, the cells were stained with 50 nM TMRM (Invitrogen, #T668), 100 nM MitoTracker Green FM (Invitrogen, #M7514), and 1 drop/mL NucBlue Live (Invitrogen, #R37605). After 20 minutes of staining, the medium was aspirated and washed with DPBS (Gibco, #14190144). The cells were then incubated in phenol red-free growth media.

During imaging, cells were kept under 5% CO_2_ and 37 °C. We used a spinning disk confocal microscope at the UCSD Nikon Imaging Center. The microscope is equipped with a spinning disk scanner (X-Light V3, CrestOptics), and an ORCA-Fusion digital CMOS camera (Hamamtsu). We used a water-immersion Plan Apo VC 60x WI NA 1.2 objective. We used 3 laser lines (405 nm, 477 nm, 546 nm) generated with a Lumencor Celesta light engine. To automate the data acquisition for multiple conditions, we used the JOBS high-content screening module on NIS Element software. For each well, the optimal plane of focus was first determined by a two-pass autofocus with a range of 80 microns. Inside each well, the software generated a random path consisting of 25 fields of view (FOVs). For each FOV visited, we performed another single-pass autofocus with a range of 20 microns to account for focus drift due to cell and plate variations in z-direction. We then captured three channels simultaneously using triggered acquisition. After all 25 FOVs were imaged, the next well was automatically visited and the protocol was repeated until all 25 drugs + 1 control wells were imaged.

### Drug Treatments

Each drug was incubated for 30 minutes prior to imaging. Drugs were present during staining and imaging. All 25 drugs were treated and imaged during one experiment, so the total incubation time ranged from 1-3 hours.

|  |  |  |  |  |
| --- | --- | --- | --- | --- |
| 1 | 2 | 3 | 4 | 5 |
| **Oligomycin** | **Valinomycin** | **Azide** | **Nocodazole** | **Antimycin A** |
| 10 µM | 100 nM | 1 mM | 10 µM | 10 µM |
| 6 | 7 | 8 | 9 | 10 |
| **Thapsigargin** | **Resveratrol** | **Rotenone** | **Ciliobrevin D** | **Nigericin** |
| 10 µM | 10 µM | 10 µM | 10 µM | 1 µM |
| 11 | 12 | 13 | 14 | 15 |
| **Cytochalasin** | **DNP** | **CCCP** | **Mitomycin C** | **Paraquat** |
| 10 µM | 10 µM | 10 µM | 10 µM | 100 µM |
| 16 | 17 | 18 | 19 | 20 |
| **MYLS22** | **MitoQ** | **Mdivi-1** | **Lonidamine** | **TBHP** |
| 10 µM | 10 µM | 10 µM | 10 µM | 1 mM |
| 21 | 22 | 23 | 24 | 25 |
| **Cyclosporine** | **P110** | **Tiron** | **Cisplatin** | **MFI8** |
| 10 µM | 1 µM | 10 µM | 10 µM | 10 µM |

### Data Preprocessing

We start with 25 FOVs for each cell type with three-channels (NucBlue Live, MitoTracker Green, TMRM). Each FOV contains many cells. The patch sizes vary for different cell types depending on the cell size. For Cal27 we chose 256×256 patches while for HeLa cells we chose 384×384 patches to reflect the larger cell size. Note that our models are generalizable across cell size, i.e. they can be trained on different image sizes and perform inference on different cell sizes. We first compute channel-wise 98th percentile values over the dataset and threshold each image channel-wise by these values to get rid of outlier high intensity values. Next, we normalize our images patch-wise, and apply patch-wise midtone-balancing or gamma correction on each image to enhance contrast and suppress noise. To get patches centered at a single cell nucleus, we apply thresholding on the nucleus channel followed by morphological opening. Finally, we apply connected component search to identify each cell nucleus. We take the centroid of each nucleus and take a fixed size 3-channel patch around that to get our final patches centered at a single nucleus. We randomly jitter the position of each patch by a few pixels to introduce natural variability into the position of the cell in the image. We extract an average of 80 cell patches from each FOV, amounting to a dataset of anywhere between 30,000 to 80,000 cell patches for different cell types.

### Self-Supervised Deep Learning

We used self-supervised frameworks for extracting the embeddings, where each embedding is the representation of an image in a lower dimensional space. Specifically, we used SimCLR and BYOL, two methods which work on the principle of comparing embeddings of augmented views of the same sample.

SimCLR is a contrastive learning method that works on the principle that augmented patches of the same images should be close together in the embedding space. Our deep learning model is a ResNet18 backbone encoder, followed by a multi-linear perceptron (MLP) head, which together reduce each multi-channel image down to a vector of 128 dimensions. We find that ResNet18 backbones are enough to model our data, but other backbones can also be used, for example, ResNet50 for larger datasets, as well as larger embedding dimensions. For self-supervised training, we feed in stochastically augmented patches of single cells to the model (the details of the augmentations are described in the next section). The model is trained to match the embeddings of augmented versions of the same patches, while pushing semantically different patches far apart. A contrastive infoNCE loss^63^ is used for this purpose, with a temperature value fixed at 0.01. We train the model with a learning rate of 0.0003, a decay rate of 0.0001, and a batch size of 256. By learning to recognize similarities and dissimilarities between various augmented images, the model delineates the inherent factors of variation in the data without requiring any labels.

BYOL is similar to SimCLR in that it also compares augmented views of the same samples. However, unlike SimCLR, it uses two networks, the online and target network, and tries to directly align the latent space embeddings of the two views projected using the two networks via a simple mean-squared error loss. We use the same ResNet18 backbone for the online and target networks, followed by an MLP projection head that reduces dimensionality down to 128 dimensions. Unlike SimCLR, we use a mean-squared-error (MSE) loss to directly align the embeddings of the online and target. The optimizer hyperparameters like learning rate and batch size are kept the same as our SimCLR models.

For our autoencoder model, we use a hierarchical vector quantized variational autoencoder^64^ with a classifier head as in Kobayashi et al.^38^. The classifier head predicts the drug label that each cell is treated with. As opposed to proxy labels, here we train the model with the actual label itself. The model has two encoders and decoders each with 4 residual blocks, with a top embedding dimension of 32 and a bottom embedding dimension of 64. We use a dictionary size of 512 embeddings. We use MSE as the reconstruction loss and categorical cross-entropy as the classification loss. The learning rate, weight decay, and batch size are kept the same as the other methods.

Our supervised learning model is a ResNet18 model trained with a categorical cross-entropy loss function. Once again, the optimizer parameters are kept the same.

All models use an Adam optimizer and are trained on graphical processing units (GPU). The end result of our deep learning models is a latent space that maps each cell image to a lower dimensional vector. Each vector captures the relevant phenotypic properties of that cell image. The latent space acts as a search space or a map, and can be reduced to a lower dimensionality for easier visualization.

Our data augmentations are defined in the next section, and are critical to implicitly teach the model to ignore spurious factors of variation. For example, to teach the deep learning model that cell orientation is not a relevant phenotypic property, we feed in pairs of rotated cell images; the model is trained to reduce the distance between them in embedding space.

### Data Augmentation

We design our augmentations to make the model specifically robust to experimental variation such as noise or cell shape. All augmentations are stochastic and are sequentially and randomly added to each individual image.

- To make the model robust to imaging noise, we add Gaussian noise of the same order as seen in the original images. Noise is added randomly to 50% of the patches with varying mean and a standard deviation which is fixed to be a particular fraction of the maximum value in each image. The mean of the additive Gaussian noise is drawn from a normal distribution centered at the mean background value of patches over the entire training set, while the variance is set at 2% to 10% of the maximum value of each patch, channel-wise.
- To make the model robust to slight variations in intensity caused by factors like imaging variation and dye uptake in cells, we use additive brightness augmentation, where we randomly add or subtract a value drawn from a Gaussian distribution with a fixed mean and variance to each image.
- To make the model robust to cell shape, orientation, and other spurious structural factors, we use random horizontal and vertical flips, random crops, random rotation, and patch-reordering (In patch reordering, we split the image into 4 patches and randomly jumble up the patches). Once again, these augmentations are only applied to a portion of images chosen randomly.

### Visualizing and Understanding the Phenotypic Latent Space

The feature representations hence created from the self-supervised method described above are 128 dimensional vectors. For visualization purposes, we reduce the dimensionality of this feature representation to 3 dimensions by using UMAP^59^. UMAP (Uniform Manifold Approximation and Projection) is a dimensionality reduction algorithm which uses a graph based approach to represent data and transform high dimensional data into lower dimensions.

The data points in UMAP are denoted by nodes in the graph and the distances or similarities between these nodes is depicted by the edges. It tries to reduce the dimensionality of the space while preserving the global structure of original data. The UMAP algorithm is computationally efficient and can handle large datasets with millions of data points. It has been widely used in data visualization, clustering, and machine learning applications.

For gaining a semantic understanding of the phenotypic latent space, we perform Principal Component Analysis to delineate the major axes of variation. We find that ∼10 principal components are enough to explain ∼75% of the variance in the embedding space. We also compute the Pearson correlation coefficient of each of these axes with cell-level mitochondrial features extracted using Mitochondria Analyzer.

## ACKNOWLEDGEMENTS

The authors thank Pham Vo and Silvio Gutkind for helpful discussions and early feedback on the manuscript and for the Cal27 cell lines. The authors thank Heng Xu for helpful discussions and his insights into PCA analysis of the latent space. The authors thank Frank Noé, Jin Zhang, and Christopher Obara for helpful suggestions during manuscript writing. This study was supported by funds from the Hartwell Foundation through an Individual Biomedical Research Award to J.S., through an NIH Director’s New Innovator Award to J.S. and through an W.M. Keck Award to J. S.

## AUTHOR CONTRIBUTIONS

### Authors and Affiliations

All authors were part of the Department of Pharmacology, University of California, San Diego, San Diego, CA, 92093 and the Department of Chemistry and Biochemistry, University of California, San Diego, San Diego, CA, 92093.

### Contributions

Parth Natekar and Johannes Schöneberg initiated the project. Parth Natekar and Mehul Arora created the deep learning based latent space, performed other deep learning-based experiments, and trained the self-supervised models. Zichen Wang was responsible for the cell-culture and microscopy. Hiroyuki Hakozaki was responsible for microscopy training. Johannes Schöneberg, Parth Natekar, Mehul Arora, and Zichen Wang wrote the manuscript. Johannes Schöneberg was responsible for conceptualization, funding, and administration.

